# Development of Rosaceae Crop-Specific Nanopore Models for Community Use Through the Genome Database for Rosaceae

**DOI:** 10.1101/2025.06.23.660393

**Authors:** Christopher Gottschalk, Erik Burchard, Lauren Whitt, Cheryl Vann, Chris Dardick, Sook Jung, Doreen Main, Jessica Marie Waite, Loren Honaas, Alex Harkess

**Affiliations:** USDA-ARS, Appalachian Fruit Research Station, Kearneysville, WV; Department of Horticulture, Washington State University, Pullman, WA; USDA-ARS, Tree Fruit Research Laboratory, Wenatchee, WA; HudsonAlpha Institute for Biotechnology, Huntsville, AL

**Keywords:** genome sequencing, basecalling models, long-read sequencing, bioinformatics, genome assembly, genotyping

## Abstract

Oxford Nanopore Technologies (ONT) sequencing platforms have enabled scientists to generate sequence reads of 20 kilobase or more, which has facilitated the assembly of complete genomes. Nanopore sequencing involves recording the change in electrical signals as the DNA molecule transverses the pore. Converting this signal information into bases – a process called basecalling – is challenging and require the implementation of machine learning. The models developed for ONT basecallers have continually improved. However, the mean quality of the reads generated fall below a Phred score of 20 (99% accuracy), the target quality for high-quality genome assembly, when used on challenging plant samples. These low-quality base calls limit the ability to conduct *de novo* genome assembly, accurate phasing of haplotypes, and long-read genotyping. To overcome these shortcomings, we fine-tuned ONT’s high accuracy basecalling model using Bonito, an AI-based deep learning basecaller, to develop crop-specific models for five important species in the Rosaceae family. These species include highly valuable tree crops such as apple, peach, pear, plum, and sweet cherry. Through the development of an automated training pipeline, we were able to achieve ∼14% higher mean and median basecall quality, and a ∼20% increase in total reads with Phred scores >15. These results were achieved without significantly affecting the read length N50s and total reads called. As a result, these new crop-specific models will enable members of the Rosaceae genomics community to produce higher quality ONT sequencing data. Furthermore, we are releasing our automated pipeline to facilitate others to train their own organism or crop-specific models.

## 1 Introduction

Third-generation sequencing technologies such as HiFi from Pacific Biosciences (Menlo Park, CA) and Oxford Nanopore Technologies (ONT; Oxford, U.K.) have democratized the sequencing and assembly of reference-quality genomes (Deamer et al., 2016; Jain et al., 2016; Wenger et al., 2019). Specifically, ONT offers a unique value proposition to scientists as the sequencing platform is relatively inexpensive and can be purchased and operated outside of sequencing centers. Moreover, ONT’s unique ability to sequence fragments ranging from kilobases to megabases has created opportunities to completely assemble genomes, such as the telomere-to-telomere (T2T) human genome (Payne et al., 2019; Amarasinghe et al., 2020; Nurk et al., 2022). To conduct ONT sequencing you must first obtain a high-molecular weight DNA sample. This DNA sample is then used to prepare a sequencing library using a kit supplied by ONT. The library is then loaded onto a flowcell and sequenced. The sequencing and standard data analysis (basecalling, filtering, demultiplexing) is conducted within a graphical user interface program called MinKnow. MinKnow and standalone basecalling software, such as ONT-provided Guppy and Dorado, convert the raw signal information into nucleotide sequences through the use of neural networks (Wick et al., 2019; Amarasinghe et al., 2020).ONT regularly releases improvements on these processes, such as new versions of library preparation kits, flow cells, and software. One area that has had significant uplift in the quality of ONT generated data is the basecalling models used by the basecalling software (Wick et al., 2019; Vereecke et al., 2020; Ferguson et al., 2022).

Currently, most studies rely on the use of ONT-released basecalling models for their data analysis. However, several studies have found that either training species-specific or fine-tuned ONT models offer increases in the Q score of the called sequences (Wick et al., 2019; Vereecke et al., 2020; Ferguson et al., 2022). These species-specific models are even more valuable for plant samples due to a lack of plant data used in training of ONT models (Amarasinghe et al., 2020; Schmidt et al.). Ferguson et al. (2020) reported gains of more than 6% for plant species when using a species-specific model compared to the ONT model with the recent flow cells (R10.4). Rosaceae is considered one of the six most important plant families with tens of domesticated crops spread throughout in member Genera (Bennett, undated). Due to this importance, we set out to develop crop specific basecalling models for five of its crops.

These crop-specific models were trained on ONT generated data using the latest library preparation kits (V14), flow cells (R10.4.1), and base callers (Bonito and Dorado). The development of these resources will be provided the Rosaceae community through their respective database, GDR (Jung et al., 2019).

## 2 Methods

### 2.1 Plant DNAs

For the *Malus* samples, leaves were collected from representative trees at the Clarksville Research Station of Michigan State University. For the *Pyrus* samples, leaves were collected from representative trees at USDA-ARS Appalachian Fruit Research Station in Kearneysville, WV, or from tissue cultured plantlets from the USDA-ARS Tree Fruit Research Laboratory in Wenatchee, WA. Leaves were flash frozen and ground into a fine powder. High-molecular weight (HMW) DNA was extracted using a Qiagen Genomic-tip 100/G kit (Germantown, MD) using a protocol developed by Driguez et al. (2021). For the *Prunus* samples (*Prunus avium, Prunus domestica,* and *Prunus persica*), leaf material was collected from representative trees located at the USDA-ARS Appalachian Fruit Research Station. HMW DNA from these three species were extracted using Dodecyltrimethylammonium bromide (DTAB) extraction protocol. Each sample had 100 mg of flash frozen leaf tissue ground in a mortar and pestle until a fine powder was obtained. The powder was transferred to a 2 mL microfuge tube and 0.6 mL of DTAB buffer was added. Each DTAB solution was incubated for 30 mins at 68°C with intermittent vortexing. Following the incubation period, 600 µL of chloroform was added to the sample and gently mixed. The sample was centrifuged for 10 mins at 16,000 g. The supernatant was pipetted off and put into a fresh microfuge tube. An equal volume of isopropanol was added and mixed, and centrifuged for 15 mis at 10,000 g. The supernatant was pipetted off without disturbing the pellet. 300 µL TE buffer was added to the microfuge tube and flicked to resuspend the DNA pellet. We then added 100 µL LiCl (8M) to final concentration of 2M and mixed gently. The TE-LiCl-DNA solution was then incubated on ice for at least two hours then centrifuged for 10 mins as 10,000 g. The supernatant was again pipetted off the pellet without disturbing it, and place into a fresh microfuge tube. We then added 2.5 volumes of EtOH to the tube with supernatant and mixed gently by inversion. A final centrifugation for 10 mins as 10,000 g was performed. The supernatant was removed by pipetting leave a pellet of DNA. The DNA pellet was then washed using 200 µL of EtOH, and centrifuged for two mins at 10,000 g. The EtOH was removed and the pellet allowed to air dried and finally resuspended in DNase free water. All extracted HMW DNA samples were checked for quantity and quality on an Agilent TapeStation (Santa Clara, CA).

### 2.2 Library Preparation and Sequencing

Sequencing libraries for each sample were prepared using a Ligation Sequencing Kit V14 (SQK-LSK114) from ONT. The preparations followed the standard protocol as provided by the kit. For *Malus domestica* ‘Cripps Pink’, we additionally prepared an ultra-long sequencing library using the Ultra-Long Kit V14 (LSK-ULK114) from ONT. Again, we followed the standard protocol as provided by the kit. Libraries were checked for quality and fragment size using an Agilent TapeStation to determine loading concentrations for sequencing. All libraries were sequenced on their own respective PromethION R10.4.1 flow cell (FLO-PRO114M). The sequencing was conducted on a PromethION P2-Solo connected to a linux workstation running MinKnow (versions varied). PromethION was operated using the default parameters.

### 2.3 Basecalling Model Training

We developed a custom bash script to automate the process. The first step of the automated workflow is indexing a reference genome (**Fig. 1**). For each species, a reference genome corresponding to the sequenced was used (**Table 1**) (Verde et al., 2017; Linsmith et al., 2019; Callahan et al., 2021; Li et al., 2024). Each reference genome was first download and short contigs (non-pseudomolecules/chromosomes) were removed. If phased haplotypes were available for the reference genomes, those were concatenated into a single FASTA file and ensured that no duplicate header lines were present. We indexed each reference genome using Minimap2 (v2.28) with default parameters (Li, 2021). We then randomly selected 250-260 POD5 files (https://github.com/nanoporetech/pod5-file-format) from the POD5 passed directory generated by each sequencing run, and placed them into separate training subdirectory, containing sets of 25 POD5 files each. The target was 1.5M reads at minimal for training spread across the POD5 files. The next step involved looping through each training subdirectory and running the ONT pytorch basecaller Bonito. Bonito (v0.8.1; https://github.com/nanoporetech/bonito) was executed with the –save-CTC and alignment function with the reference genome index as input with the --reference parameter. Since we aimed to fine-tune, Bonito was also executed using the ONT reference dna_r10.4.1_e8.2_400bps_hac@v5.0.0 basecalling model. The third step was to then merge the resulting numpy chunk, reference lengths, and reference arrays from each executed Bonito run using a custom python script. The fourth step was to fine-tune the ONT reference dna_r10.4.1_e8.2_400bps_hac@v5.0.0 basecalling model using Bonito train function. Here, we executed Bonito with --epochs 5, --lr 5e-4, --pretrained dna_r10.4.1_e8.2_400bps_hac@v5.0.0 and the directory containing the merged chunk, reference lengths, and reference arrays as input. The output from Bonito train is a fine-tuned basecalling model. The fifth and final step used Bonito export on the resulting fine-tune model to generate a Dorado compatible fine-tune model (**Fig. 1**).

**Figure 1.**
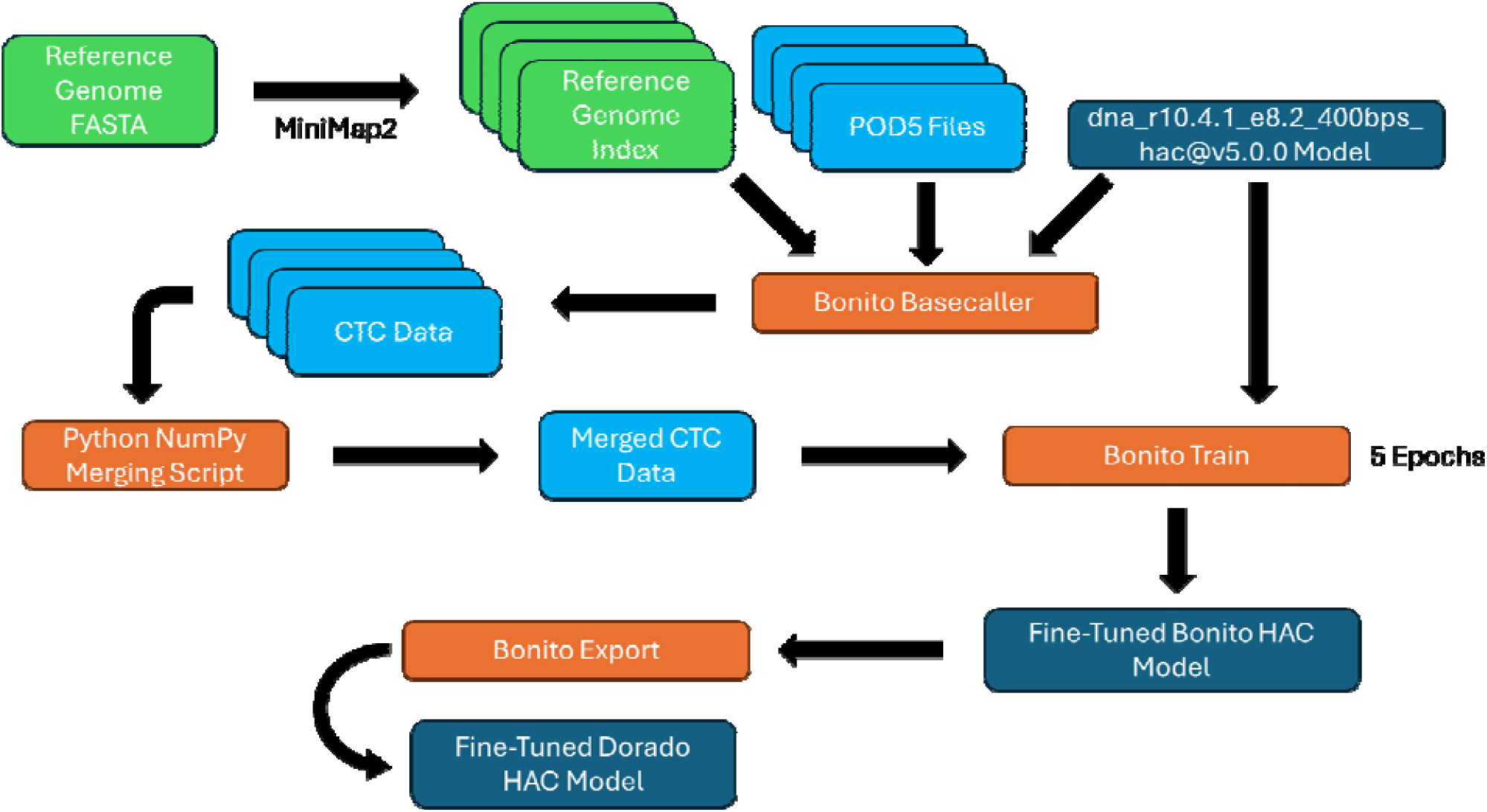
Graphical illustration of the training of fine-tuned basecalling models.

**Table 1.**
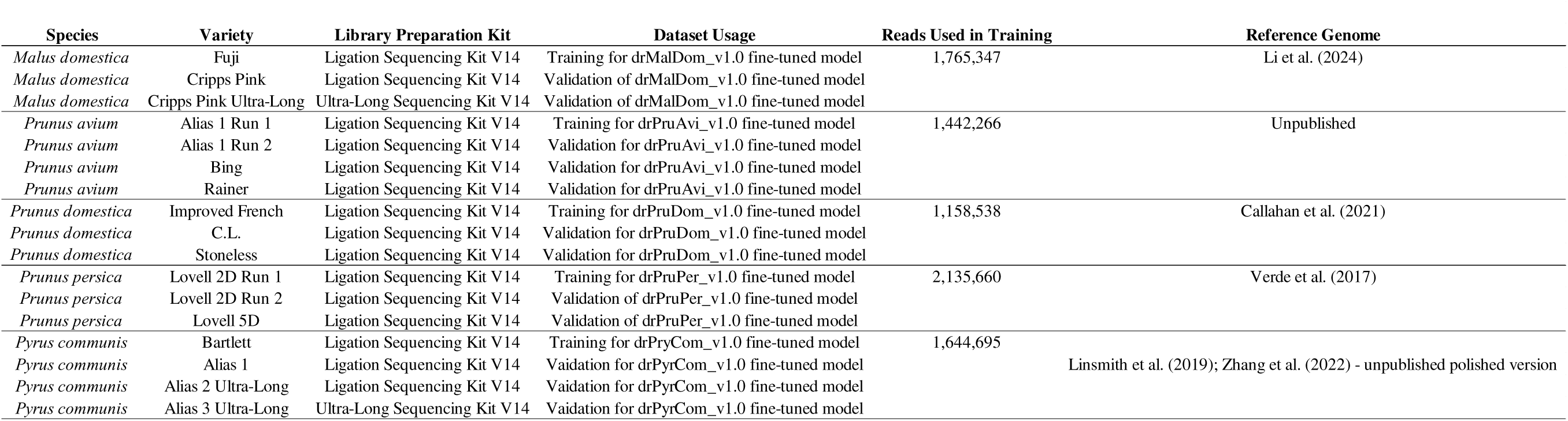
Sequencing datasets, library preparations information, dataset usage, reads used for training, and corresponding reference genomes used.

### 2.4 Fine-Tuned Model Evaluation

To evaluate the performance of the fine-tuned models for each crop we utilized the Dorado (v.0.9.0; https://github.com/nanoporetech/dorado) basecaller. Here, the entire POD5 dataset for each sample used for training and additional sequencing of other varieties were basecalled with Dorado using the ONT reference dna_r10.4.1_e8.2_400bps_hac@v5.0.0 basecalling model and again with the newly trained fined-tuned model (**Table 1**). The resulting basecalled output BAM files were converted to FASTQ using SAMTools fastq (v.1.17; Li et al., 2009; Danecek et al., 2021). The FASTQ files were then filtered for reads with Q scores <9, if not already performed during sequencing, by using Chopper (v.0.9.2; De Coster & Rademakers, 2023). We then used the filtered reads as input into NanoPlot and NanoComp to obtain statistics and graphical representations of the data (De Coster & Rademakers, 2023).

### 2.5 Data Availability

The custom scripts used in the training of the fine tune models are available in a GitHub Repository https://github.com/gottsc33/ONT_FineTuner. The resulting fine-tuned models for each crop have been released through genome database/repositories for each crop. For example, Rosaceae crops (apple, pear, and *Prunus*) are available on the Genome Database for Rosaceae (GDR) (Jung et al., 2019). The raw POD5 files used in training and validation can be made available upon appropriate request.

## 3 Results and Discussion

We generated sixteen sequencing datasets that span five crops in the Rosaceae family (**Table 1 and 2**). Five of the sixteen sequencing datasets were selected for use in training of crop-specific basecalling models, while the remaining eleven were saved for validation purposes. The five sequencing datasets selected for training of crop-specific models were chosen due to the availability of previously published genomes (e.g. ‘Fujì apple) or an internal unpublished genome that was available and is being released in this study (**Table 1**). Our sequencing generated between a minimum of 7.42 Gb and a maximum of 87 Gb, and a total of 518.17 Gb (**Table 2**). Fifteen of the datasets contained over 1M reads with N50 ranging from 10 - 12 Kb for some *Prunus* samples to 43 Kb for a *Pyrus* sample generated using the Ultra-Long Library Preparation kit from ONT. In total, we generated over 66 M reads with a read N50 estimated to be 22 Kb (**Table 2**). We generated sixteen sequencing datasets that span five crops in the Rosaceae family (**Table 1 and 2**). Five of the sixteen sequencing datasets were selected for use in training of crop-specific basecalling models while the remaining eleven were saved for validation purposes. The five sequencing datasets selected for training of crop-specific models were chosen based on the availability of previously published genomes (e.g. ‘Fujì apple) or an internal unpublished genome that was available and is being released in this study (**Table 1**)(Verde et al., 2017; Linsmith et al., 2019; Callahan et al., 2021; Li et al., 2024). Our sequencing generated between a minimum of 7.42 Gb and a maximum of 87 Gb, and a total of 518.17 Gb (**Table 2**). Fifteen of the datasets contained over 1M reads with N50 ranging from 10 - 12 Kb for some *Prunus* samples to 43 Kb for a *Pyrus* sample generated using the Ultra-Long Library Preparation kit from ONT. In total, we generated over 66 M reads with a read N50 estimated to be 22 Kb (**Table 2**).

**Table 2.**
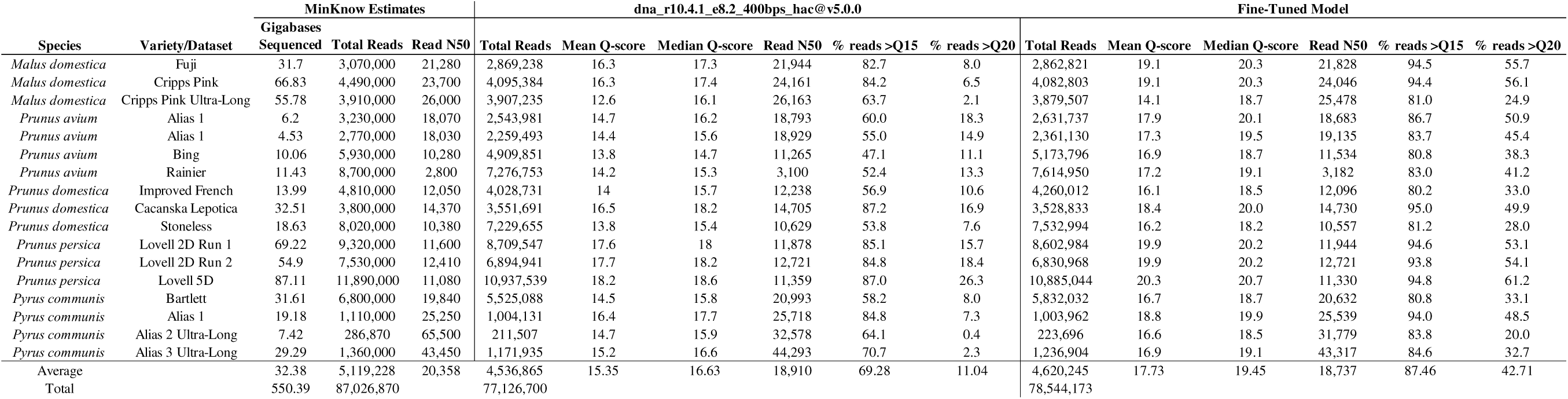
Comparison between the dna_r10.4.1_e8.2_400bps_hac@v5.0.0 and fine-tuned crop models.

For apple (*Malus domestica*), we trained a crop-specific model using ‘Fujì resequencing data, utilizing a recently published reference genome for that variety (Li et al., 2024). Following training on over 1.7 M reads, our new apple base calling model (drMalDom_v1.0_dna_r10.4.1_e8.2_400bps_hac@v5.0.0) demonstrated improved performance compared to the ONT reference model (dna_r10.4.1_e8.2_400bps_hac@v5.0.0) (**Fig. 2**, **Table 1 and 2**). This trend was consistent for basecalling on the full ‘Fujì dataset and two ‘Cripps Pink’ datasets that were not used in the training of the model (**Table 1 and 2**). For example, we observed significant increases in the mean and median Q-scores and the percentage of reads >Q15 (**Table 2** and **3**). These increases were calculated to be between 16 and 17% (**Table 3**). We also observed small decreases in the number of total reads called and the read N50 lengths at <2% change. However, neither of those two metrics were found to be significantly lower when comparing the two different models. The percentage of reads >Q20 exhibited a significant increase of 823% when using the fine-tuned model, which represents the largest percentage change between the two models (**Table 2** and **3**).

**Figure 2.**
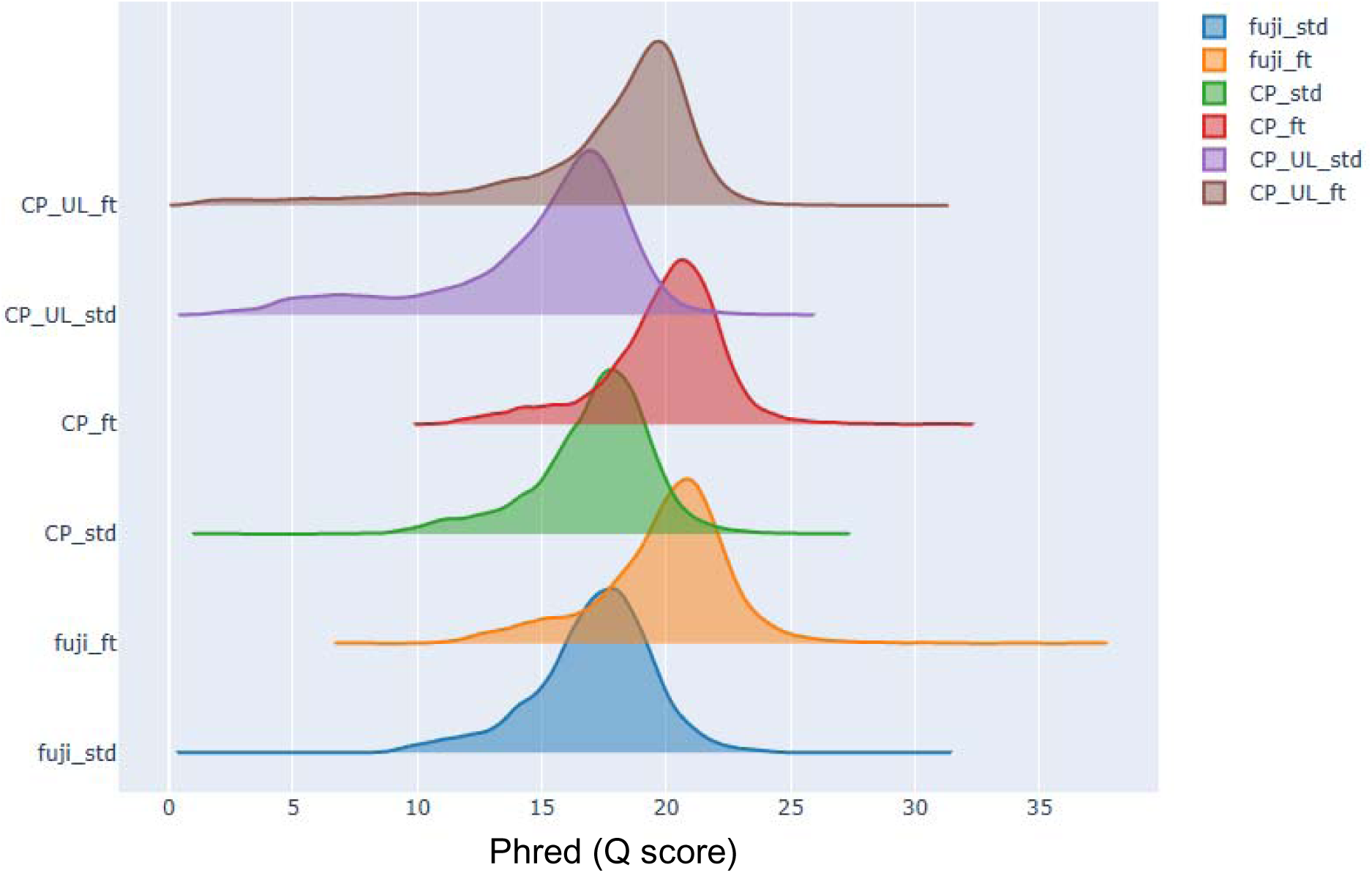
Ridge plot of the distribution of phred scores in the paired *Malus* sequencing datasets. Std – standard nanopore model; ft – fine-tuned apple model; UL – ultra-long; CP –’Cripps Pink.

**Table 3.**
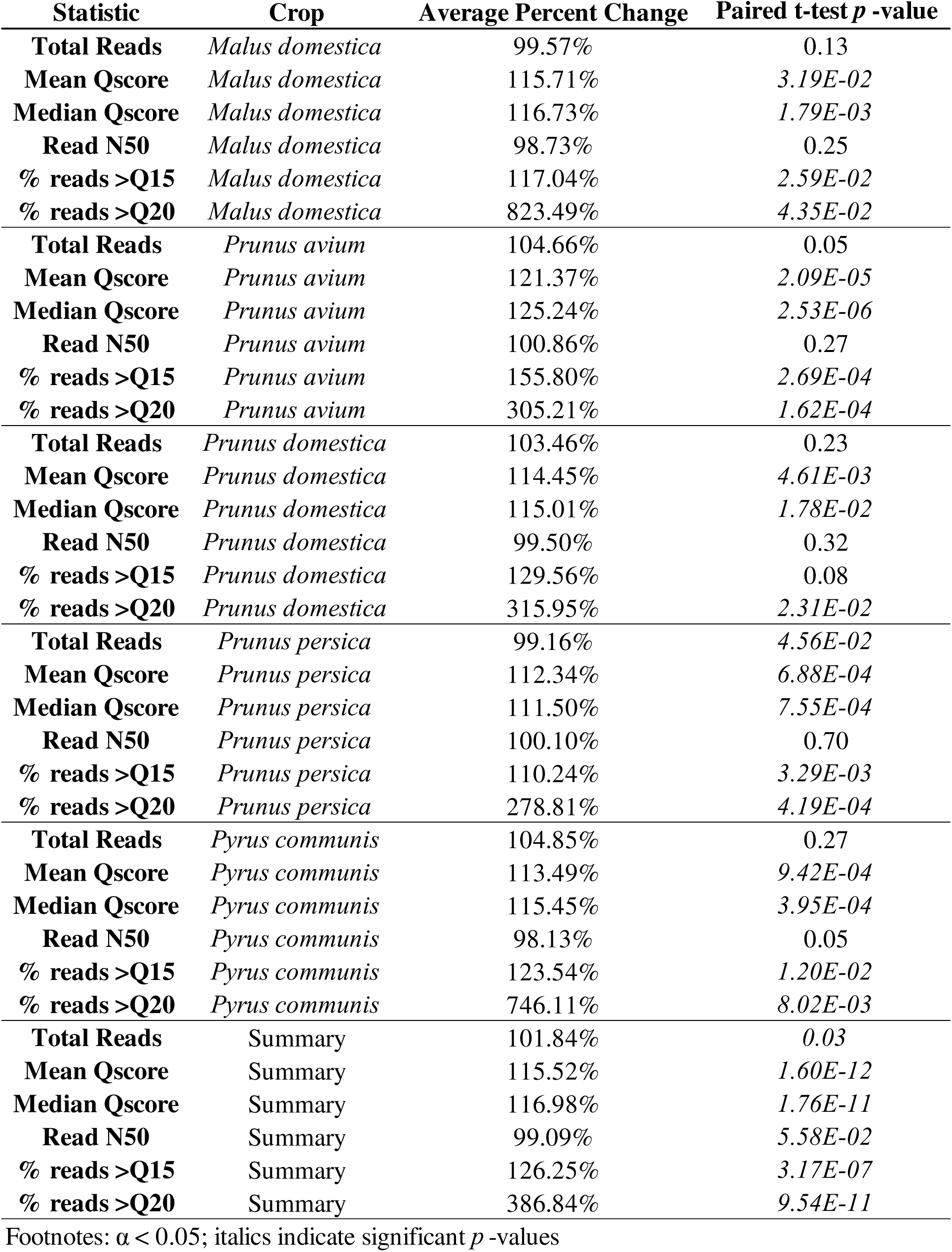
Statistical tests on the differences between the usage of the fine-tuned vs. ONT reference basecalling model.

For sweet cherry (*Prunus avium*), we trained crop-specific models using an undisclosed variety (*P. avium* A1) sequence data. We had access to an unpublished reference genome assembled for this variety. Following training on over 1.4 M reads, our new sweet cherry base calling model (drPruAvi_v1.0_dna_r10.4.1_e8.2_400bps_hac@v5.0.0) demonstrated a similar and significant increase in performance over the ONT reference model in the mean and median Q-scores and the percentage of reads >Q15 (**Fig.3A**, **Table 2** and **3**). These increases were calculated to be between 21 and 55% (**Table 3**). We also observed a small but significant increase in the number of total reads called (**Table 3**). The read N50 was virtually the same, with a small 0.86% increase using the fine-tuned model. However, this metric was not found to be significant (**Table 2** and **3**). The percentage of reads >Q20 exhibited a significant increase of 305% when using the fine-tuned model, which represents the largest percentage change between the two models (**Table 2** and **3**).

**Figure 3.**
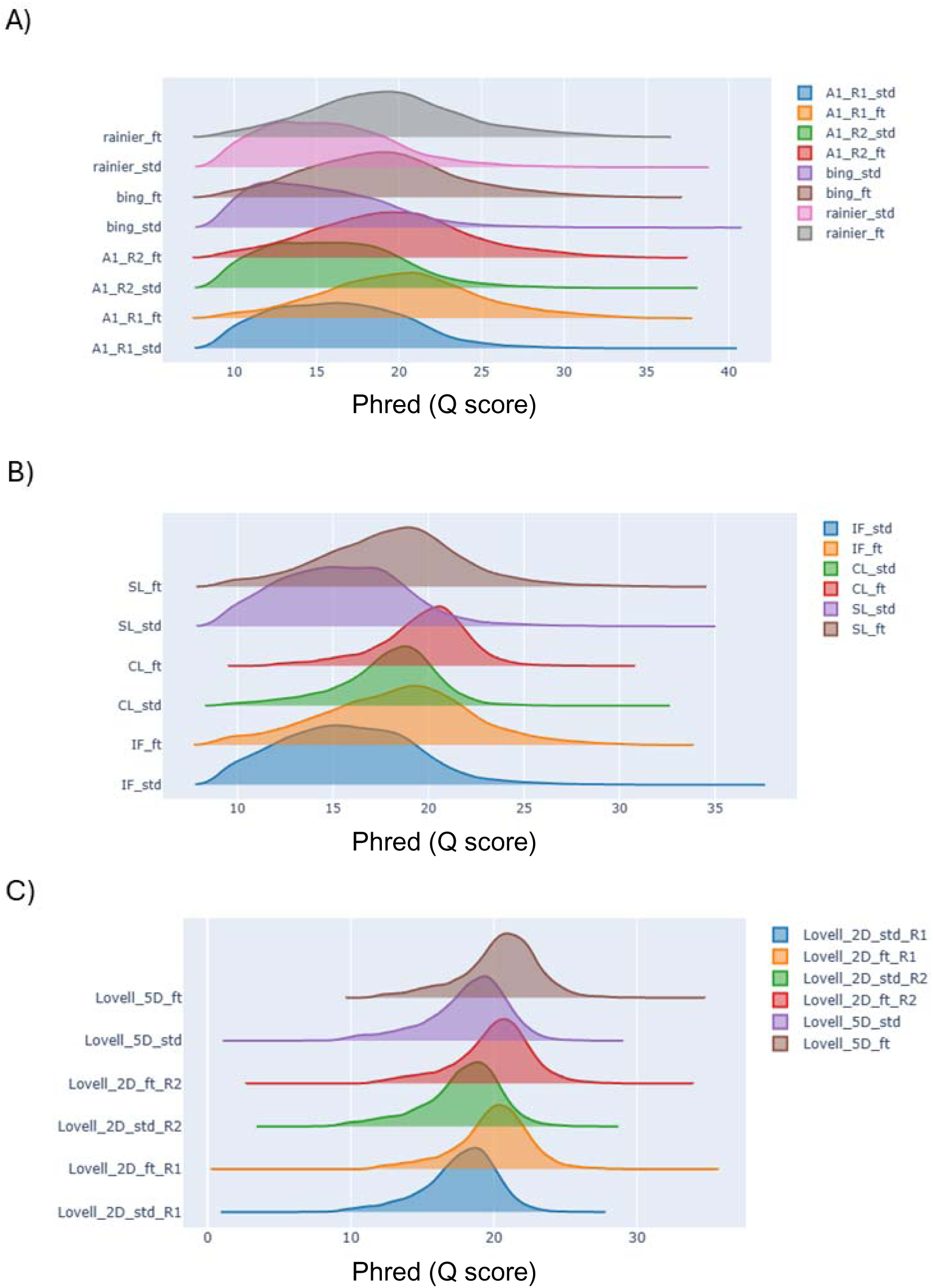
Ridge plot of the distribution of phred scores in the paired *Prunus* sequencing datasets. A) *P. avium,* B) *P. domestica*, and C) *P. persica*. Std – standard nanopore model ; ft – fine-tuned apple model; UL – ultra-long; A - Alias; R – Biological replicate sequencing sample

For plum (*Prunus domestica*), we trained crop-specific models using ‘Improved French’ sequence data. A previously published reference genome for this variety was available (Callahan et al. 2021). Following training on over 1.1 M reads, our new plum base calling model (drPruDom_v1.0_dna_r10.4.1_e8.2_400bps_hac@v5.0.0) demonstrated a similar and significant increase in performance over the ONT reference model in the mean and median Q-scores and the percentage of reads >Q15 (**Fig. 3B**, **Table 2** and **3**). These increases were calculated to be between 14 and 30% (**Table 3**). The total number of reads and read length N50 were nearly the same, with a small 3% increase using the fine-tuned model for total reads and <1% decrease in read N50. These two metrics were not found to be significant (**Table 2** and **3**). The percentage of reads >Q20 exhibited a significant increase of 315% when using the fine-tuned model, which represents the largest percentage change between the two models (**Table 2** and **3**). This result is similar to what was observed with sweet cherry but lower than in apple.

For peach (*Prunus persica*), we trained crop-specific models using ‘Lovell 2D’resequence data. *P. persica* had a reference genome already available for that variety (Verde et al., 2013, 2017). Following training on over 2.1M reads, our new peach base calling model (drPruPer_v1.0_dna_r10.4.1_e8.2_400bps_hac@v5.0.0) demonstrated a similar and significant increase in performance over the ONT reference model in the mean and median Q-scores and the percentage of reads >Q15 (**Fig. 3C**, **Table 2** and **3**). These increases were calculated to be between 10 and 12% (**Table 3**). We also observed a small but significant decrease in the number of total reads called (**Table 3**). The read length N50 was virtually the same, with a small 0.1% increase using the fine-tuned model. However, this metric was not found to be significant (**Table 2** and **3**). The percentage of reads >Q20 exhibited a significant increase of 278% when using the fine-tuned model, which represents the largest percentage change between the two models (**Table 2** and **3**). This result is again similar to what was observed with the other two *Prunus* species.

For pear (*Pyrus communis*), we trained crop-specific models using ‘Bartlett’ sequence data. *P. communis* has a doubled haploid genome available (Linsmith et al. 2019), however, we instead used an in-house polished version of this ‘Bartlett’ doubled haploid genome that is unpublished (**Table 1**)(Zhang et al., 2022). Following training on 1.6 M reads, we observed a similar increase in performance as all the other data sets using the crop-specific model in the mean and median Q-scores and the percentage of reads >Q15 (**Fig. 4**, **Table 2** and **3**). Again, these increases were calculated to be between 13 and 23% (**Table 3**). Similar, lower and not significant read N50 were found in the crop-specific model datasets. Similar to the *P. persica* datasets, total reads were increased in the fine-tuned model but was not significant (**Table 3**). The percentage of reads >Q20 exhibited a significant increase of 746% when using the fine-tuned model, which represents the largest percentage change between the two models (**Table 2** and **3**). This result is similar to what was observed with the *Malus* data sets.

**Figure 4.**
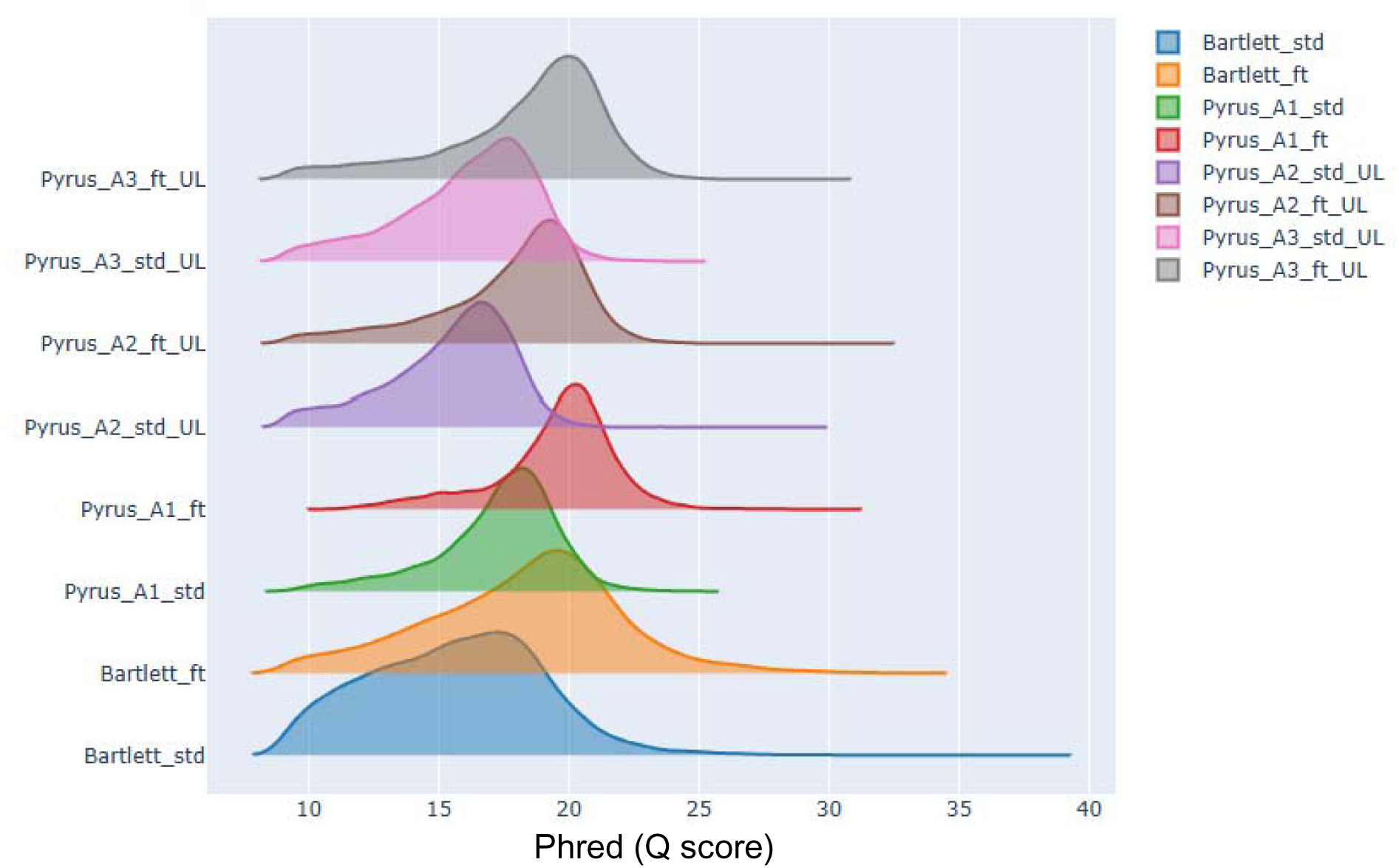
Ridge plot of the distribution of phred scores in the paired *Pyrus* sequencing datasets. Std – standard nanopore model; ft – fine-tuned apple model; UL – ultra-long; A – Alias.

To evaluate if the fine-tuned models were suitable for basecalling in an intergeneric fashion, we executed basecalling using all five of the crop specific models on three dataset that were not used in the training (**Table 4**). These datasets included representatives from *P. avium, M. domestica,* and *P. communis.* Consistently across all three experiments the model developed for that species performed the best across all Q-score metrics (**Table 4**). For example, in the *M. domestica* experiment the drMalDom_v1.0 (apple) model had the highest mean and median Q-scores and output the most reads >Q15 and >Q20. The closely related drPyrCom_v1.0 (pear) model was the second best and subsequently followed by drPruPer_v1.0 (peach), drPruAvi_v1.0 (sweet cherry), and drPruDom_v1.0 (plum). This order does not hold true for the other experiments but in all cases the fine-tune crops specific models outperformed the reference model (dna_r10.4.1_e8.2_400bps_hac@v5.0.0) from ONT. These results suggest that within family (e.g. Rosaceae) basecalling can be performed with any of these models and achieve higher quality data over the reference model.

**Table 4.**
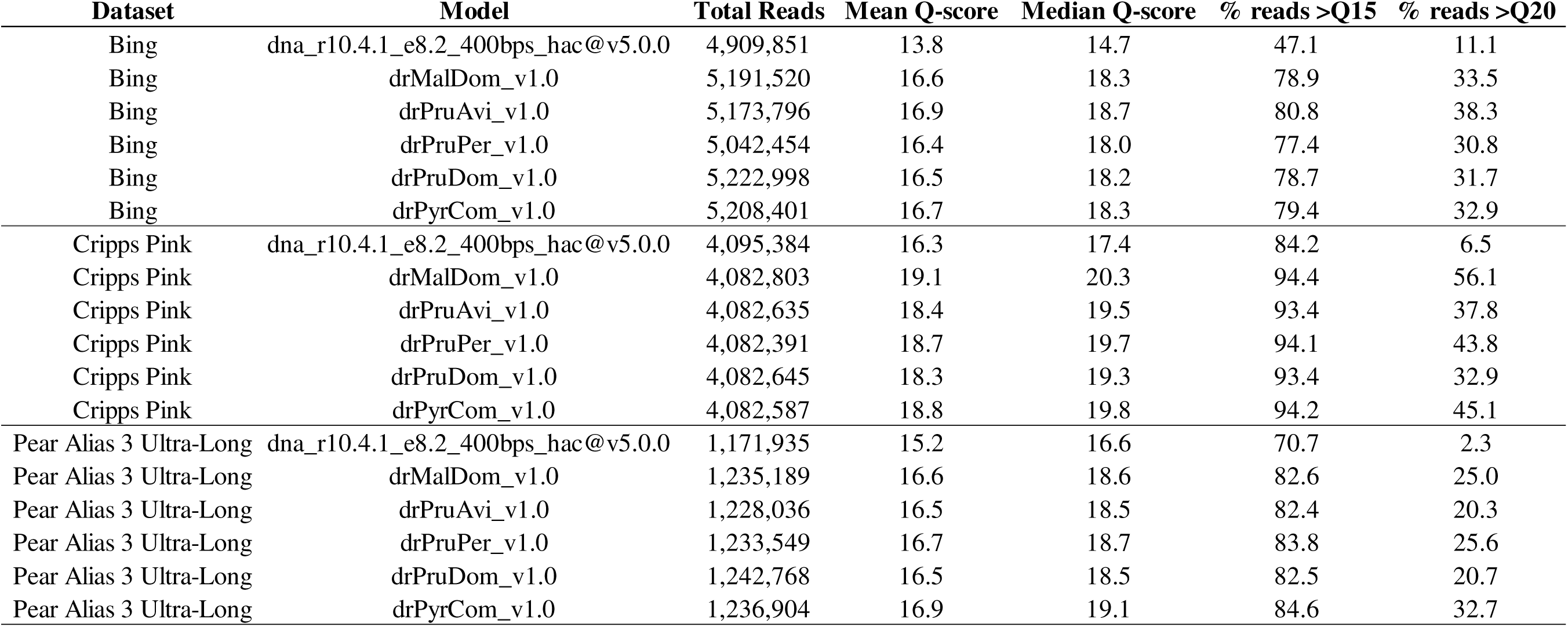
Comparing the read quality of different crop specific fine-tuned basecalling models applied to intergeneric datasets.

## 4 Conclusions

Overall, our fine-tuning of Oxford Nanopore’s dna_r10.4.1_e8.2_400bps_hac@v5.0.0 model for crop specific increase the mean and median Q scores and increased the dataset’s percent of reads >Q15 and >Q20 to significant levels (**Table 3**). These increases were around the 14% improvement in the Q score metrics and 19% increase in the available reads >Q15. However, the average read N50 was decreased by a significant 1.26% in the fine-tuned models (**Table 3**). The tradeoff between a slight decrease in read N50 vs overall quality is still favorable for genome assembly and genotyping purposes. The metric that exhibited the greatest increase in improvement was the percentage of reads >Q20 (**Table 3**). Here, for/using the entire data set of 17 sequencing runs, we observed a significant 386% average increase in the number of reads with >Q20 scores. The training of these initial crop specific models for Rosaceae crops will help advance the research community and promote the use of in-house sequencing methods. These models were only developed for the high-accuracy model class. This class is a middle group between rapid but error-prone basecalling (fast models) and the super accurate (sup) models. However, the pipeline developed in this work can be used to fine-tune these additional model classes. Moreover, we demonstrated that the crop specific models can also be used in an intergeneric fashion within the Rosaceae family and outperform the ONT reference model.

## Acknowledgements

We would like to thank Pairwise for allowing access to an unpublished high-quality *P. avium* genome for training. Additionally, we would like to thank Dr. Courtney Hollender and Alex Englesma for providing the tissue samples for ‘Fujì and ‘Cripps Pink’ apples. We additionally want to thank Pairwise (Durham, NC) for providing unpublished sequences used in this manuscript.

## Author Contributions

CG conceptualized the study, organized, tested, and wrote the pipeline scripts, sample collection and conduction of research, and wrote the original draft of the manuscript. Study design input was also obtained by CD. Sample collection, research, and dataset availability were provided by CV, EB, AE, CH, JMW, LH, AY, and AH. The release and management of the model resources was conducted by DM and SJ. All authors had input on manuscript content, conducted editing and review, and approved final manuscript.

## References

Amarasinghe, S.L., Su, S., Dong, X., Zappia, L., Ritchie, M.E., & Gouil, Q. (2020). Opportunities and challenges in long-read sequencing data analysis. Genome Biology, 21, 30. 10.1186/s13059-020-1935-5

Bennett, B.C. (undated). Twenty-Five Economically Important Plant Families. In B.C. Bennett (Ed.), Economic Botany Florida International University, Miami, USA.

Callahan, A.M., Zhebentyayeva, T.N., Humann, J.L., Saski, C.A., Galimba, K.D., Georgi, L.L., Scorza, R., Main, D., & Dardick, C.D. (2021). Defining the ‘HoneySweet’ insertion event utilizing NextGen sequencing and a de novo genome assembly of plum (Prunus domestica). Horticulture Research, 8, 8. 10.1038/s41438-020-00438-2

Danecek, P., Bonfield, J.K., Liddle, J., Marshall, J., Ohan, V., Pollard, M.O., Whitwham, A., Keane, T., McCarthy, S.A., Davies, R.M., & Li, H. (2021). Twelve years of SAMtools and BCFtools. GigaScience, 10, giab008. 10.1093/gigascience/giab008

De Coster, W., & Rademakers, R. (2023). NanoPack2: population-scale evaluation of long-read sequencing data. Bioinformatics, 39, btad311. 10.1093/bioinformatics/btad311

Deamer, D., Akeson, M., & Branton, D. (2016). Three decades of nanopore sequencing. Nature Biotechnology, 34, 518–524. 10.1038/nbt.3423

Ferguson, S., McLay, T., Andrew, R.L., Bruhl, J.J., Schwessinger, B., Borevitz, J., & Jones, A. (2022). Species-specific basecallers improve actual accuracy of nanopore sequencing in plants. Plant Methods, 18, 1–11. 10.1186/s13007-022-00971-2

Jain, M., Olsen, H.E., Paten, B., & Akeson, M. (2016). The Oxford Nanopore MinION: delivery of nanopore sequencing to the genomics community. Genome Biology, 17, 239. 10.1186/s13059-016-1103-0

Jung, S., Lee, T., Cheng, C.-H., Buble, K., Zheng, P., Yu, J., Humann, J., Ficklin, S.P., Gasic, K., Scott, K., Frank, M., Ru, S., Hough, H., Evans, K., Peace, C., Olmstead, M., DeVetter, L.W., McFerson, J., Coe, M., Wegrzyn, J.L., Staton, M.E., Abbott, A.G., & Main, D. (2019). 15 years of GDR: New data and functionality in the Genome Database for Rosaceae. Nucleic Acids Research, 47, D1137–D1145. 10.1093/nar/gky1000

Li, H. (2021). New strategies to improve minimap2 alignment accuracy. Bioinformatics, 37, 4572–4574. 10.1093/bioinformatics/btab705

Li, H., Handsaker, B., Wysoker, A., Fennell, T., Ruan, J., Homer, N., Marth, G., Abecasis, G., Durbin, R., & 1000 Genome Project Data Processing Subgroup. (2009). The Sequence Alignment/Map format and SAMtools. Bioinformatics, 25, 2078–2079. 10.1093/bioinformatics/btp352

Li, W., Chu, C., Li, H., Zhang, H., Sun, H., Wang, S., Wang, Z., Li, Y., Foster, T.M., López-Girona, E., Yu, J., Li, Y., Ma, Y., Zhang, K., Han, Y., Zhou, B., Fan, X., Xiong, Y., Deng, C.H., Wang, Y., Xu, X., & Han, Z. (2024). Near-gapless and haplotype-resolved apple genomes provide insights into the genetic basis of rootstock-induced dwarfing. Nature Genetics, 56, 505–516. 10.1038/s41588-024-01657-2

Linsmith, G., Rombauts, S., Montanari, S., Deng, C.H., Celton, J.-M., Guérif, P., Liu, C., Lohaus, R., Zurn, J.D., Cestaro, A., Bassil, N.V., Bakker, L.V., Schijlen, E., Gardiner, S.E., Lespinasse, Y., Durel, C.-E., Velasco, R., Neale, D.B., Chagné, D., Van de Peer, Y., Troggio, M., & Bianco, L. (2019). Pseudo-chromosome-length genome assembly of a double haploid “Bartlett” pear (Pyrus communis L.). GigaScience, 8, giz138. 10.1093/gigascience/giz138

Nurk, S., Koren, S., Rhie, A., Rautiainen, M., Bzikadze, A.V., Mikheenko, A., Vollger, M.R., Altemose, N., Uralsky, L., Gershman, A., Aganezov, S., Hoyt, S.J., Diekhans, M., Logsdon, G.A., Alonge, M., Antonarakis, S.E., Borchers, M., Bouffard, G.G., Brooks, S.Y., Caldas, G.V., Chen, N.-C., Cheng, H., Chin, C.-S., Chow, W., Lima, L.G. de, Dishuck, P.C., Durbin, R., Dvorkina, T., Fiddes, I.T., Formenti, G., Fulton, R.S., Fungtammasan, A., Garrison, E., Grady, P.G.S., Graves-Lindsay, T.A., Hall, I.M., Hansen, N.F., Hartley, G.A., Haukness, M., Howe, K., Hunkapiller, M.W., Jain, C., Jain, M., Jarvis, E.D., Kerpedjiev, P., Kirsche, M., Kolmogorov, M., Korlach, J., Kremitzki, M., Li, H., Maduro, V.V., Marschall, T., McCartney, A.M., McDaniel, J., Miller, D.E., Mullikin, J.C., Myers, E.W., Olson, N.D., Paten, B., Peluso, P., Pevzner, P.A., Porubsky, D., Potapova, T., Rogaev, E.I., Rosenfeld, J.A., Salzberg, S.L., Schneider, V.A., Sedlazeck, F.J., Shafin, K., Shew, C.J., Shumate, A., Sims, Y., Smit, A.F.A., Soto, D.C., Sović, I., Storer, J.M., Streets, A., Sullivan, B.A., Thibaud-Nissen, F., Torrance, J., Wagner, J., Walenz, B.P., Wenger, A., Wood, J.M.D., Xiao, C., Yan, S.M., Young, A.C., Zarate, S., Surti, U., McCoy, R.C., Dennis, M.Y., Alexandrov, I.A., Gerton, J.L., O’Neill, R.J., Timp, W., Zook, J.M., Schatz, M.C., Eichler, E.E., Miga, K.H., & Phillippy, A.M. (2022). The complete sequence of a human genome. Science, . 10.1126/science.abj6987

Payne, A., Holmes, N., Rakyan, V., & Loose, M. (2019). BulkVis: a graphical viewer for Oxford nanopore bulk FAST5 files 35, 2193–2198

Schmidt, M.H.-W., Vogel, A., Denton, A.K., Istace, B., Wormit, A., van de Geest, H., Bolger, M.E., Alseekh, S., Maß, J., Pfaff, C., Schurr, U., Chetelat, R., Maumus, F., Aury, J.-M., Koren, S., Fernie, A.R., Zamir, D., Bolger, A.M., & Usadel, B. De Novo Assembly of a New Solanum pennellii Accession Using Nanopore Sequencing

Verde, I., Abbott, A.G., Scalabrin, S., Jung, S., Shu, S., Marroni, F., Zhebentyayeva, T., Dettori, M.T., Grimwood, J., Cattonaro, F., Zuccolo, A., Rossini, L., Jenkins, J., Vendramin, E., Meisel, L.A., Decroocq, V., Sosinski, B., Prochnik, S., Mitros, T., Policriti, A., Cipriani, G., Dondini, L., Ficklin, S., Goodstein, D.M., Xuan, P., Fabbro, C.D., Aramini, V., Copetti, D., Gonzalez, S., Horner, D.S., Falchi, R., Lucas, S., Mica, E., Maldonado, J., Lazzari, B., Bielenberg, D., Pirona, R., Miculan, M., Barakat, A., Testolin, R., Stella, A., Tartarini, S., Tonutti, P., Arús, P., Orellana, A., Wells, C., Main, D., Vizzotto, G., Silva, H., Salamini, F., Schmutz, J., Morgante, M., & Rokhsar, D.S. (2013). The high-quality draft genome of peach (Prunus persica) identifies unique patterns of genetic diversity, domestication and genome evolution. Nature Genetics, 45, 487–494. 10.1038/ng.2586

Verde, I., Jenkins, J., Dondini, L., Micali, S., Pagliarani, G., Vendramin, E., Paris, R., Aramini, V., Gazza, L., Rossini, L., Bassi, D., Troggio, M., Shu, S., Grimwood, J., Tartarini, S., Dettori, M.T., & Schmutz, J. (2017). The Peach v2.0 release: high-resolution linkage mapping and deep resequencing improve chromosome-scale assembly and contiguity. BMC Genomics, 18, 225. 10.1186/s12864-017-3606-9

Vereecke, N., Bokma, J., Haesebrouck, F., Nauwynck, H., Boyen, F., Pardon, B., & Theuns, S. (2020). High quality genome assemblies of Mycoplasma bovis using a taxon-specific Bonito basecaller for MinION and Flongle long-read nanopore sequencing. BMC Bioinformatics, 21, 517. 10.1186/s12859-020-03856-0

Wenger, A.M., Peluso, P., Rowell, W.J., Chang, P.-C., Hall, R.J., Concepcion, G.T., Ebler, J., Fungtammasan, A., Kolesnikov, A., Olson, N.D., Töpfer, A., Alonge, M., Mahmoud, M., Qian, Y., Chin, C.-S., Phillippy, A.M., Schatz, M.C., Myers, G., DePristo, M.A., Ruan, J., Marschall, T., Sedlazeck, F.J., Zook, J.M., Li, H., Koren, S., Carroll, A., Rank, D.R., & Hunkapiller, M.W. (2019). Accurate circular consensus long-read sequencing improves variant detection and assembly of a human genome. Nature Biotechnology, 37, 1155– 1162. 10.1038/s41587-019-0217-9

Wick, R.R., Judd, L.M., & Holt, K.E. (2019). Performance of neural network basecalling tools for Oxford Nanopore sequencing. Genome Biology, 20, 129. 10.1186/s13059-019-1727-y

Zhang, H., Wafula, E.K., Eilers, J., Harkess, A.E., Ralph, P.E., Timilsena, P.R., dePamphilis, C.W., Waite, J.M., & Honaas, L.A. (2022). Building a foundation for gene family analysis in Rosaceae genomes with a novel workflow: A case study in Pyrus architecture genes. Frontiers in Plant Science, 13. 10.3389/fpls.2022.975942

